# Scientific evaluation of negative exome sequencing followed by systematic scoring of candidate genes to decipher the genetics of neurodevelopmental disorders

**DOI:** 10.1101/588517

**Authors:** Benjamin Büttner, Sonja Martin, Anja Finck, Maria Arelin, Carolin Baade-Büttner, Tobias Bartolomaeus, Peter Bauer, Astrid Bertsche, Matthias K. Bernhard, Saskia Biskup, Nataliya Di Donato, Magdeldin Elgizouli, Roland Ewald, Constanze Heine, Yorck Hellenbroich, Julia Hentschel, Sabine Hoffjan, Susanne Horn, Frauke Hornemann, Dagmar Huhle, Susanne B. Kamphausen, Wieland Kiess, Ilona Krey, Alma Kuechler, Ben Liesfeld, Andreas Merkenschlager, Diana Mitter, Petra Muschke, Roland Pfäffle, Tilman Polster, Ina Schanze, Jan-Ulrich Schlump, Steffen Syrbe, Dagmar Wieczorek, Martin Zenker, Johannes R. Lemke, Diana Le Duc, Konrad Platzer, Rami Abou Jamra

## Abstract

**Background:** Deciphering the monogenetic causes of neurodevelopmental disorders (NDD) is an important milestone to offer personalized care. But the plausibility of reported candidate genes in exome studies often remains unclear, which slows down progress in the field.

**Methods:** We performed exome sequencing (ES) in 198 cases of NDD. Cases that remained unresolved (n=135) were re-investigated in a research setting. We established a candidate scoring system (CaSc) based on 12 different parameters reflecting variant and gene attributes as well as current literature to rank and prioritize candidate genes.

**Results:** In this cohort, we identified 158 candidate variants in 148 genes with CaSc ranging from 2 to 11.7. Only considering the top 15% of candidates, 14 genes were already published or funneled into promising validation studies.

**Conclusions:** We promote that in an approach of case by case re-evaluation of primarily negative ES, systematic and standardized scoring of candidate genes can and should be applied. This simple framework enables better comparison, prioritization, and communication of candidate genes within the scientific community. This would represent an enormous benefit if applied to the tens of thousands of negative ES performed in routine diagnostics worldwide and speed up deciphering the monogenetic causes of NDD.

## Background

The entirety of rare diseases represents a substantial burden and affects millions of individuals worldwide^1^. A large part of these diseases have a genetic cause^2^, but many cases remain undiagnosed. A missing diagnosis may have a negative impact on the caring families, medical treatment, socio-economic expenses, and on prognosis^3^.

This is most obvious in the genetic etiologies of neurodevelopmental disorders (NDD). Exome sequencing (ES) has proven to be highly successful in identifying causes of NDD^4,5^. Thus, large efforts have been undertaken in order to decipher the genetics of NDD^6–12^. The DDD study is the most prominent example^12^. It included 7580 trio ES cases with NDD and 14 genes were reported as statistically valid novel NDD genes. Major studies like this are of high importance and give unprecedented insights into the genetics of NDD. Considering the available supplemental tables with a high number of de novo candidate variants, we understand that many NDD genes did not reach the significance threshold after correcting for multiple testing. Thus, we see that a complementary approach that includes the specific characteristics of the variants and the genes as well as the clinical information of the patients may overcome the burden of statistical significance thresholds. This requires a detailed case by case evaluation as presented by other research approaches that have focused on smaller, often homogeneous cohorts^13,14,15^. However, in these studies the plausibility of many of the reported candidate genes remained unclear. The challenge of differentiating relevant genes from false positives is an issue in single cases or small cohorts. As such small cohorts are available at many institutions worldwide, genes cannot be statistically validated. On the other hand, such institutions collectively perform tens of thousands of exome sequencing in NDD cases, including the implementation of case-specific information. More than half of them remain negative in a diagnostic setting^5^, representing a large potential in the re-evaluation of such cases in a research setting.

The above mentioned aspects and our own experience motivated us to this study ^5,14^. We demonstrate that re-evaluation of negative ES in a research setting, including the specific characteristics of the candidate variants and genes as well as the clinical information, followed by standardized scoring of identified variants and genes is a powerful tool to speed up deciphering the genetics of NDD.

## Materials and Methods

### Patients

We considered all NDD cases that were introduced to us between January 2016 and December 2017. We performed ES, when a genetic diagnosis could not be set based on routine methods such as chromosome, array, or panel analysis. In total, 198 cases were sequenced. In 92 % (n=182) of the cases, a trio setting were performed, while the remaining were solo (n=7), duo (n=4), or quattro exome sequencing (n=5). In 61% (n=120), the index was male, and in 12% (n=24) of consanguineous families. The range of age of the analyzed individuals was from the day of birth to 44 years; 9% (n=17) were under one year old, 47% (n=94) were 1 to 5 years old, 41% (n=82) were 6 to 20 years old and 3% (n=5) were older than 20 years. Epilepsy was reported in 53% (n=104) of the cases, 24% (n=48) had dysmorphic features, 20% (n=40) had microcephaly, 20% (n=39) had muscular hypotonia, 12% (n=23) had short stature, 10% (n=19) had features of autism spectrum disorder, and 5% (n=9) had macrocephaly (detailed clinical information in Case Overview in table S1). Human Phenotype Ontology (HPO) terms were used to standardize the descriptions of the phenotypic information^16^.

### Exome sequencing and bioinformatic analyses

ES was performed after an enrichment with Nextera All Human 37Mb or Agilent SureSelect All Human Version 6 (60Mb) and sequenced on an Illumina platform (HiSeq2500 or NovaSeq6000) at Centogene’s laboratory in Rostock, Germany, or at CeGaT’s laboratory in Tübingen, Germany. Analysis of the raw data including variant calling and annotation was performed using the software VARVIS (Limbus, Rostock).

### Variant evaluation

First, ES data was evaluated in a setting of routine diagnostics, meaning that results of genes that have been clearly associated with a specific phenotype were reported to the referring physicians. This evaluation based on common standards of impact of the variant, prevalence in the general population, segregation in the family, and overlapping of patients symptoms with the described phenotype, and the variants were classified based on the ACMG recommendations^17^.

If in the first step no convincing variant was identified, we evaluated the exome sequencing in a second step in a research setting. We identified candidate variants based on their minor allele frequency in the general population (genome aggregation database)^18^, on their impact on the protein using Sequencing Ontology terms^19^ and based on the segregation in the family (comparable to the suggested procedure by MacArthur and colleagues^20^).

### Candidate score (CaSc)

Due to the large number of candidate variants and in order to reduce arbitrariness in deciding which genes to follow on, to efficiently communicate with colleagues, and to focus resources on the most convincing candidates, we sought to prioritize in a top-down relevance list to compare plausible and highly convincing genes with less relevant genes^20^. For this purpose, we developed a candidate scoring (CaSc) system. At the same time, such scoring should not exclude genes that due to missing information show weak evidence of relevance, but that may get relevant in the future through accumulation of scientific knowledge.

CaSc comprises four major groups, containing 12 different parameters overall (Table 1, Figure 1, and Table S4 for detailed information). The maximal achievable CaSc is 15 points.

**Table 1:**
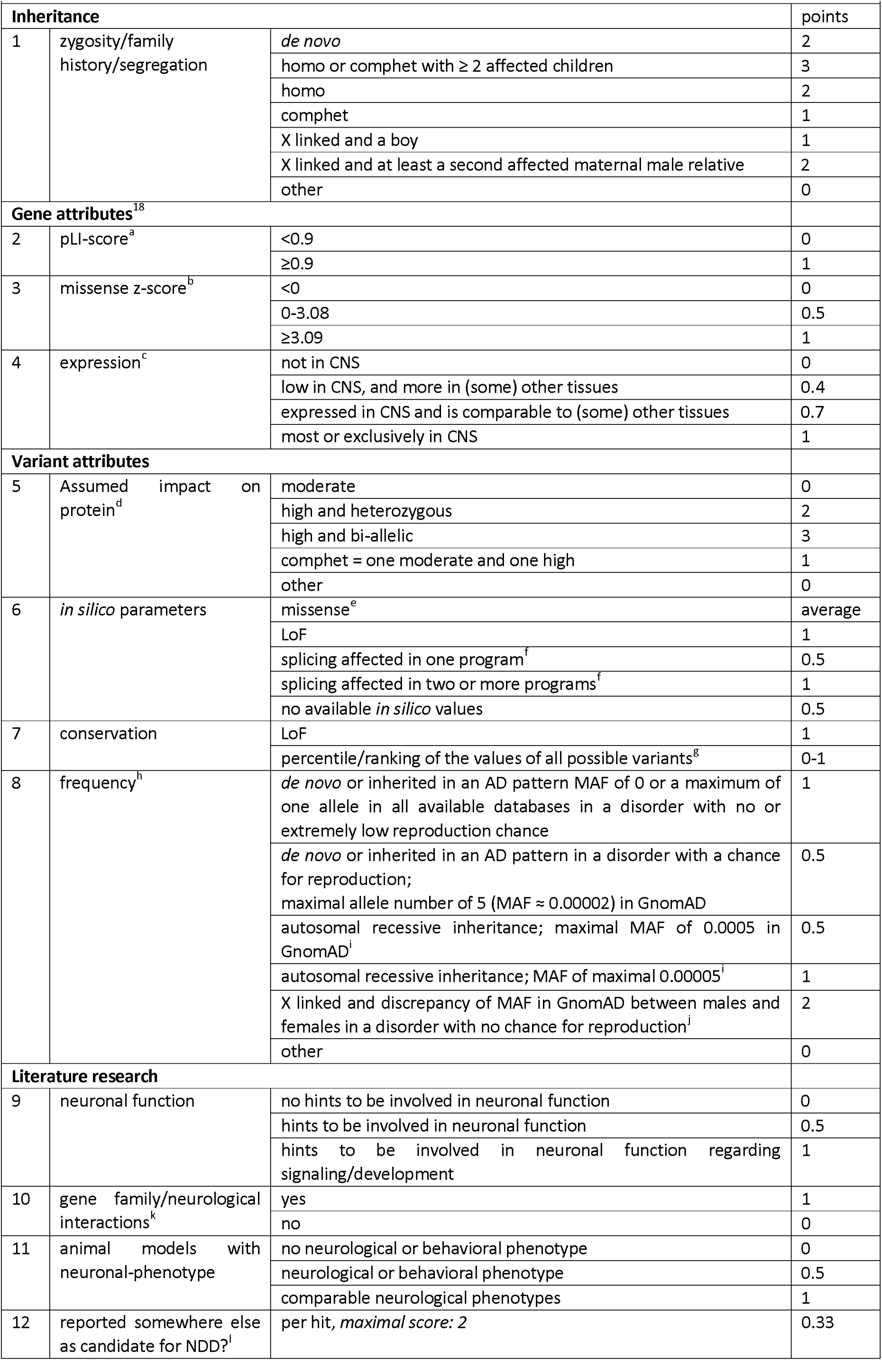
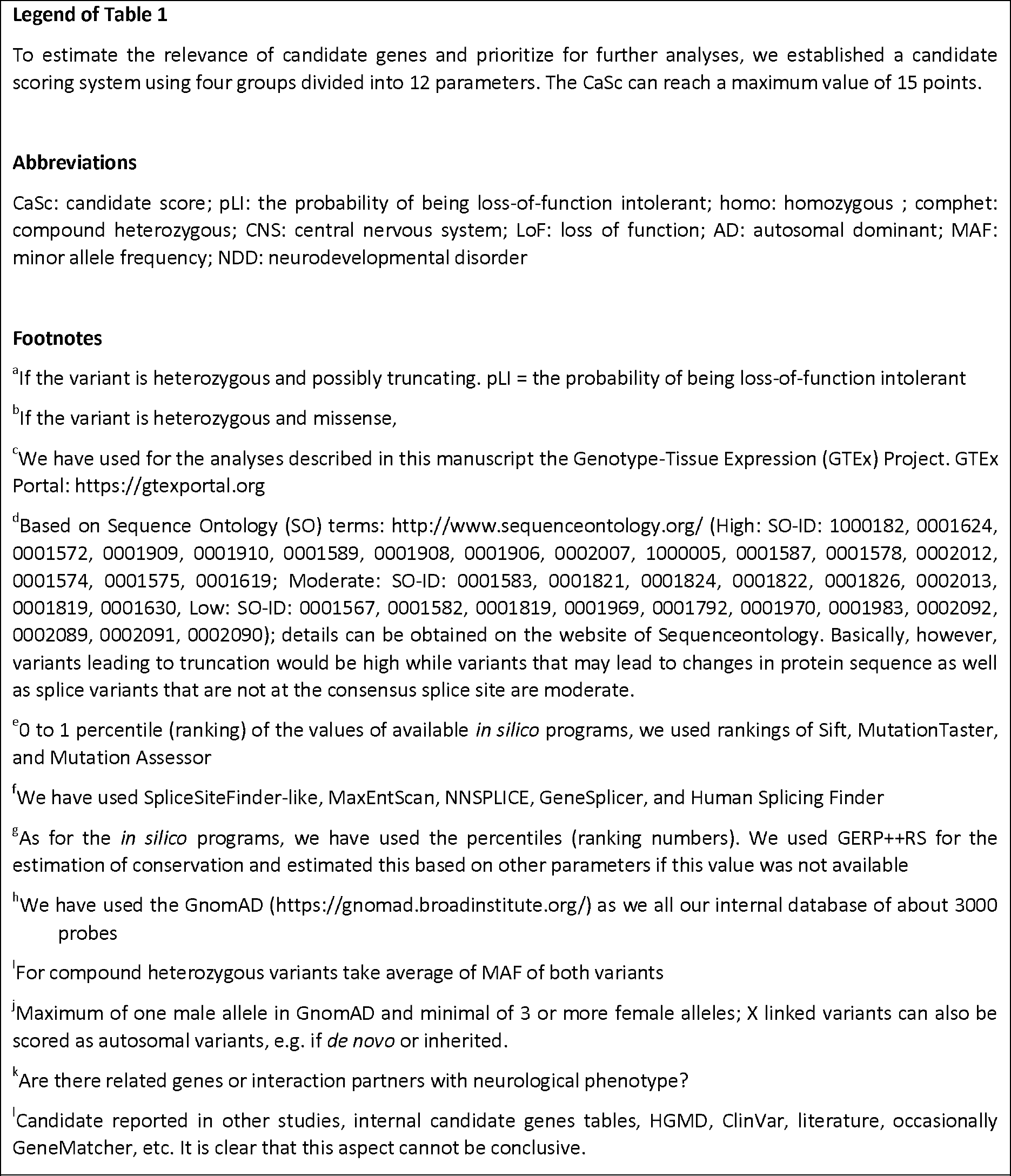
Candidate Score (CaSc)

**Figure 1:**
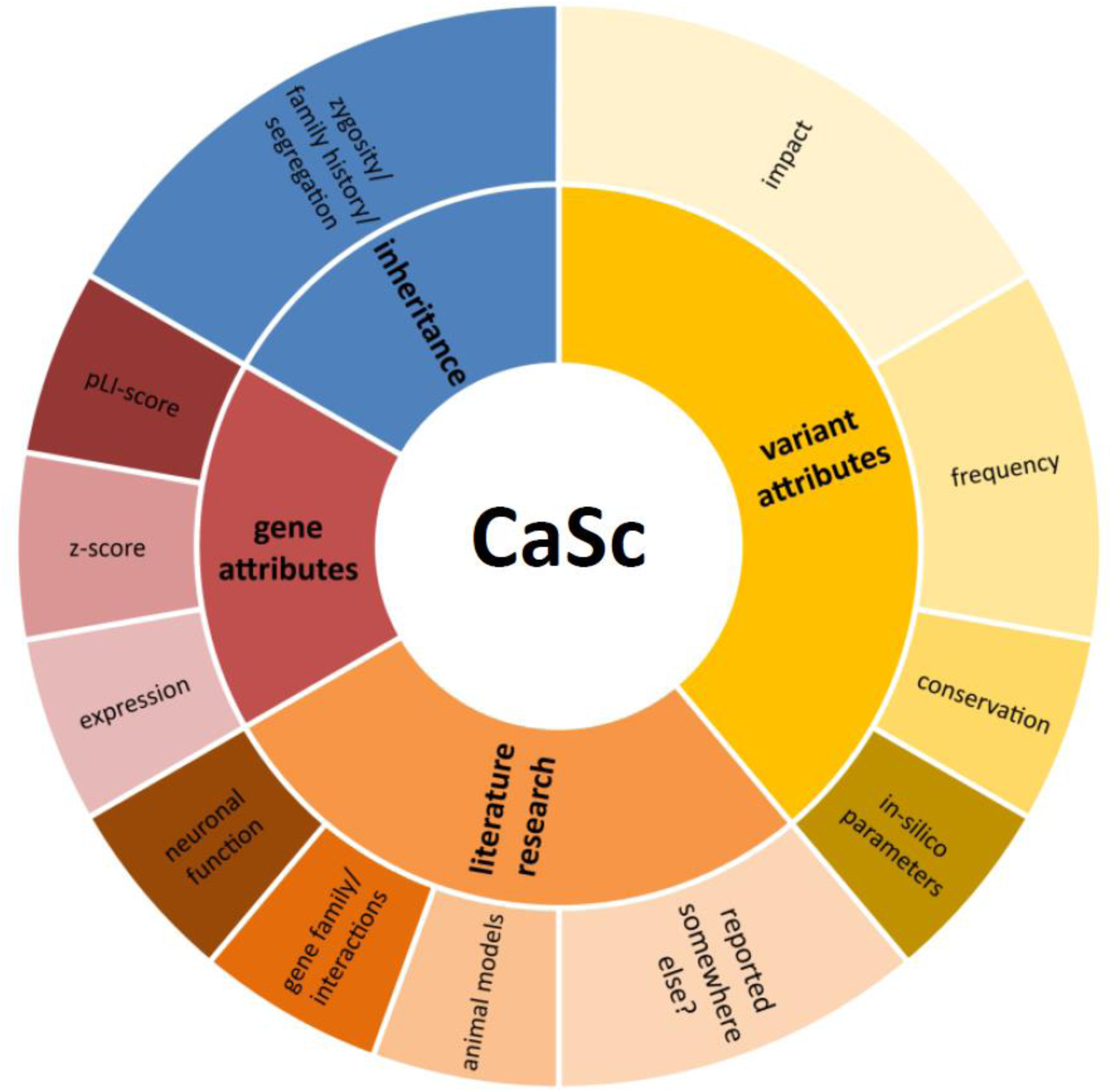
Components and structure of CaSc. CaSc comprises four major groups (in blue, red, orange and yellow color shades) containing 12 different parameters; the maximum score of each parameter varies between 1 and 3 points. The total of the points results in the CaSc (see Table 1 and Table S4 for details on the parameters and how these are evaluated)

The parameters 1, 2, 3, 5, 6, 7, and 8 of the CaSc are objective. Five parameters (4, 9, 10, 11, and 12) have a subjective component (see also Table 1 and detailed information in Table S4). This partial subjectivity is due to differences between the scientific evaluators, including individual experience in identification and assessment of relevant literature or animal models or in understanding neurological networks. On the other hand, having several parameters makes error in evaluating one of these less relevant for the sum of the CaSc, thus attenuating the subjectivity of the score (see statistic evaluation of CaSc).

### Reproducibility of CaSc

To evaluate the reproducibility of the CaSc, three different scientists scored the same 29 randomly selected candidate variants. The three scientists evaluated the subjective parameters 4, 9, 10, 11, and 12 of the CaSc (expression, neuronal function, gene family and interactions, animal model, and reported somewhere else as candidate for NDD, respectively). We performed a one-way ANOVA between the three evaluators to prove the interrater correlation. In order to exclude that an evaluator would, e.g., tend to score high genes that another scores low and vice versa, i.e. to support the specificity of the scoring, we also asked for specific scoring of each gene.

## Results

### Diagnostic yield

In 32% (n=63) of the cases we have identified variants that we found reliable enough to be reported to the referring physicians (Tables S1 and S2). Based on the ACMG-Guidelines^17^ we classified 28% of them as pathogenic, 50% as likely pathogenic, and 22% as variant of unknown significance (VUS). These cases were excluded from re-evaluation in a research setting. Details are in Figure 2 and the Supplemental Tables S1 and S2.

**Figure 2:**
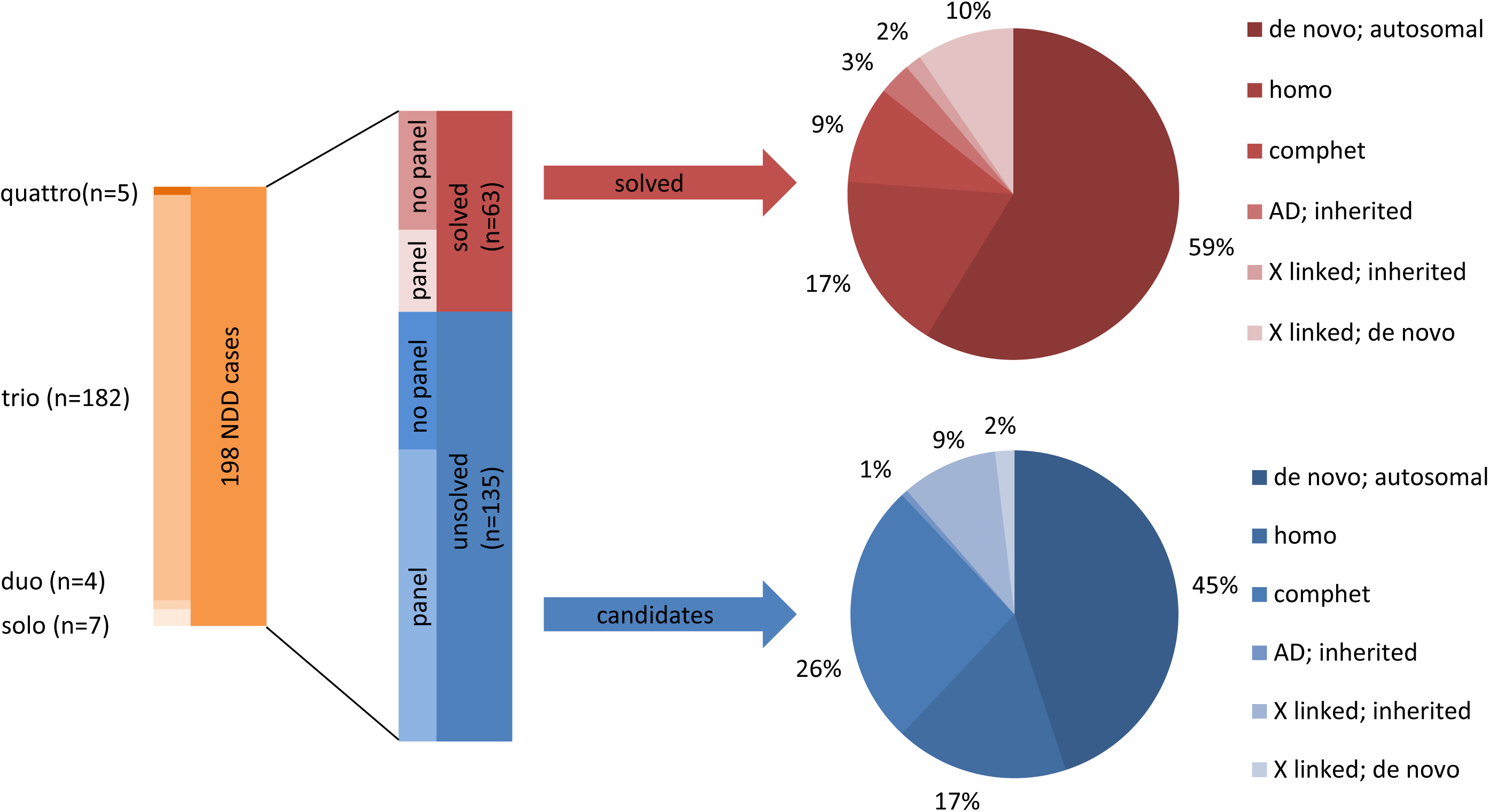
Design and yield of the NDD cohort. We performed exome sequencing of 198 NDD cases in solo, duo, trio and quattro designs (left, different orange color shades). The diagnostic yield (middle panel) varied between cases that were preceded by a panel diagnostic and that were analyzed initially in an exome setting. The right panel shows the zygosity of the diagnosed cases in the red circle and in the blue circle the zygosity of the candidate genes.

### Candidate genes

Re-evaluation of the unsolved 135 cases in a research setting revealed, in 79 cases, 158 potentially causative candidate variants in 148 candidate genes (two compound heterozygous variants count as one). CaSc varied over the 158 variants between 2.0 and 11.7 points (Figure 3). To demonstrate the utility of this score, we list below the candidates with CaSc ≥ 9 (top 15%); *TANC2, GLS, ASIC1A, KMT2E, ACTL6B, GRIN3B, SPEN, DNAH14, CUX1, PUM1, UNC13A, RASGRP1, GRIA4, SLC32A1, PUM2, TOB1, MAPK8IP3, NPTX1, ETV5, CACNB4, WDFY3* (for a detailed list of all candidates, see Table S2). Fourteen of these were published or funneled into subsequent validation studies^21–29^ (*TANC2, KMT2E*, and *WDFY3* are under review, but still unpublished data). Of the 158 candidate variants, most were *de novo* (n=74, 47%, of these 71 are autosomal, 3 are X linked), followed by autosomal recessive (n=68, 43%, of these 41 are compound heterozygous, and 27 are homozygous, mostly (n=22) observed in consanguineous families), and inherited variants (n=15, 9%, are X linked and one is an autosomal, paternally inherited heterozygous variant) (Figure 2 and Tables S1 and S2).

**Figure 3:**
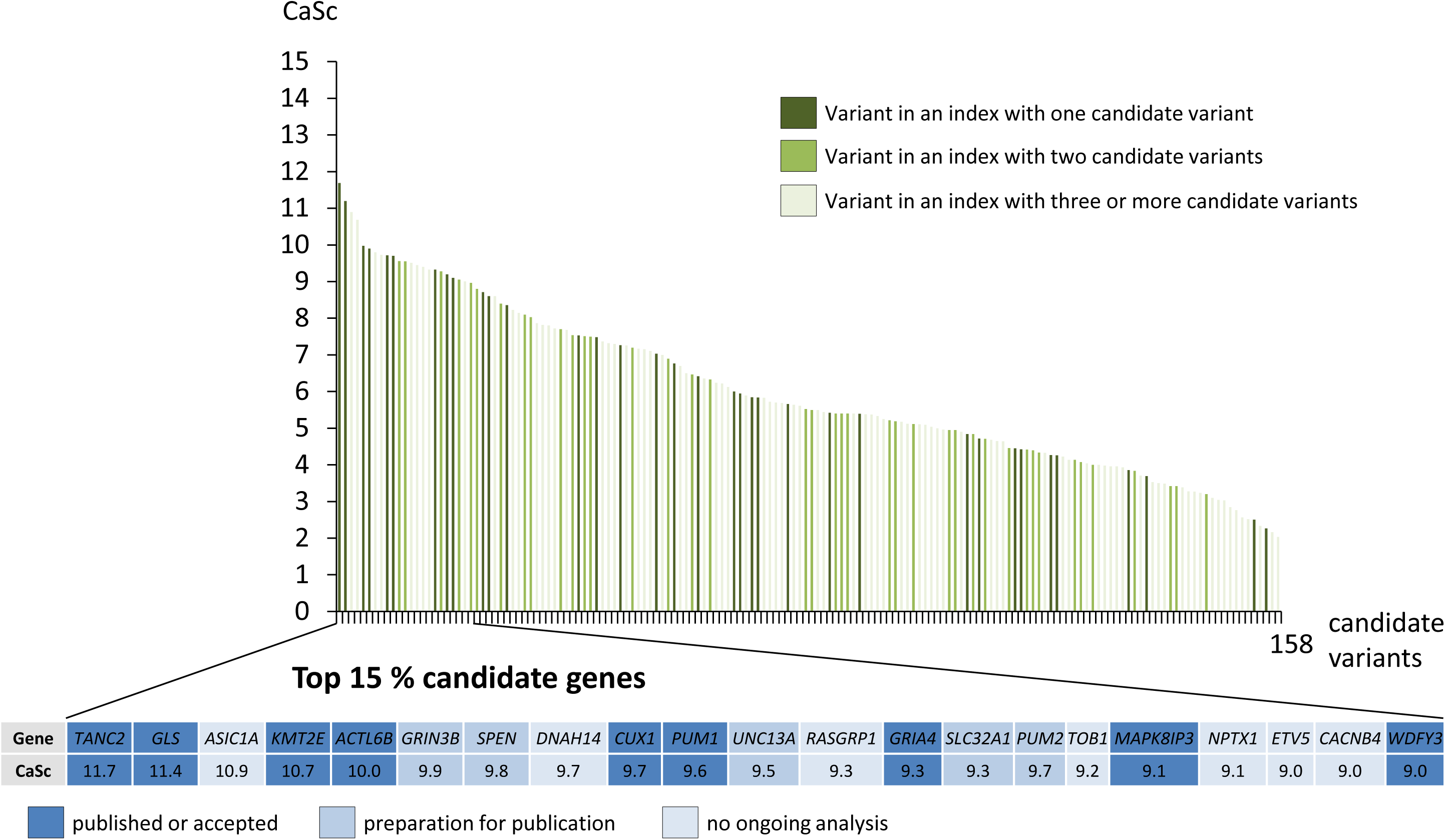
Overview and top 15% of CaSc. The maximal achievable CaSc is 15. In the upper panel we show a distribution over the 158 variants between 2.0 and 11.7 points. Different green color shades show how many candidate genes were identified in one case. The top 15% of the CaSc scored candidate genes (CaSc≥9), equivalent to 23 gene variants (and 21 genes since *GLS* and *SPEN* were hit twice). We divided these candidate genes in 3 groups; published or accepted for publication, in preparation for publication, and not yet investigated, i.e. searching for cooperation partners.

### Reproducibility of the score

To test the reproducibility of CaSc, three different users independently scored the subjective components of the same randomly selected 29 candidate variants. The CaSc of the 29 triple analyzed variants fluctuates with a mean value of 0.4 points with a standard deviation of 0.19. The one-way ANOVA showed that there is no significant difference between the evaluators (p of 0.5163). Also a pair-wise comparison of the evaluators showed no significant difference (see Table S3). To support the specificity of the scoring we also asked whether there is a difference between the scores for each gene. This demonstrates that the scoring is specific for each gene (p < 0.0001, F(28,56) = 49.71).

## Discussion

Institutions around the globe perform presumably tens of thousands of ES in cases of NDD on a yearly basis. These cases are evaluated in detail in a clinical diagnostic setting but over half of them remain negative^5^. The scientific content of these negative cases is often not fully exploited, as candidate variants and genes cannot be proven as valid on a statistical basis as demonstrated by major studies,^6–12^ due to the data residing in different “silos”. Considering the above mentioned aspects, we see the need for a systematic and complimentary approach to major NDD studies. We recommend a detailed evaluation of all negative cases, considering functional aspects of the candidate genes, *in silico* evaluation of the variants, the literature and clinical presentation, followed by a systematic and standardized scoring of candidates to prioritize genes for validation studies.

In our relatively small cohort of 198 cases with neurodevelopmental disorders, we diagnosed 63 cases due to variants in previously described NDD genes (detailed information in Figure 2). We evaluated the remaining 135 negative ES cases in a research setting. In 79 cases we identified 158 candidate variants in 148 candidate genes. It was promptly clear that many of these genes would be false positive (e.g. among cases with more than one candidate (Figure 3)). For valid disease genes, ACMG standards and guidelines are used to standardize the evaluation of variants^17^ and reduce errors. Applying feasible evaluation standards for candidate genes of NDD would ease the analysis of these variants and increase its usability for other scientists. Thus, to standardize prioritization of possible disease-causing variants, we established a candidate score (CaSc) system. This is a system of 12 parameters that can be applied to any variant within a few minutes by exome evaluators. CaSc varied in our candidates between 2 and 11.7 points (maximum range 15 points). The candidates at the lower end of CaSc have naturally a higher probability of being false positive, but including these increases the sensitivity and represents a negative control as suggested before^20^. We compensated the consequently reduced specificity by a top-down approach.

Considering all 23 candidate variants (in 21 genes) with scores of 9.0 or more (equivalent to top 15%) revealed that at least 14 of them *(TANC2, GLS, KMT2E, ACTL6B, GRIN3B, SPEN, CUX1, PUM1, UNC13A, GRIA4, SLC32A1, PUM2, MAPK8IP3, WDFY3)* are already published or funneled into projects with several other patients and often with functional analyses^21–29^ (as of 15^th^ of February 2019, see also Figure 3) (*TANC2, KMT2E*, and *WDFY3* are under review, but still unpublished data). For the remaining seven genes *ASIC1, DNAH14, RASGRP1, TOB1, NPTX1, ETV5*, and *CACNB4* there is partly good supporting evidence in the literature or via GeneMatcher^30^, but further analyses are necessary. Such genes should be followed on in the next step. The yield of genes beyond the 15% threshold is currently smaller, partially since we decided to concentrate our limited resources on genes at the top of our ranking. However, there are still some interesting genes such as *MADD* (CaSc of 7.8, top-down position: 35, manuscript in preparation) as well as *EGR3* and *GTPBP2* (CaSc of 8.7 and 8.4, respectively, and several matches in GeneMatcher). This shows that although we expect more false positive in the remaining 85% of candidates, many of these are going to be validated. Thus, we have included detailed supplemental tables of all available genetic and clinical information on each of the candidates. This offers the scientific community the possibility to find further hits for their candidates, associated with information on the relevance to enhance plausibility.

The CaSc can be roughly divided into a gene-dependent component and a variant-dependent component. The latter is fully objective and can be automatized since it includes information that is often included in a standard annotation pipelines (impact on protein, *in silico* prediction and conservation, minor allele frequency, and zygosity). The gene-dependent component is partially subjective since it includes manual evaluation of the literature regarding protein function and interactions, animal models, tissue expression, as well as identifying further hits in the gene in other studies. We are aware that the subjectivity of this information can be reduced by detailed prescription of sources and information that are allowed to be used. However, we found that this either leads to loss of information or it makes scoring a tedious task thus reducing compliance and/or efficacy of exome evaluators. In addition to the gene- and variant-components, pLI- and z-scores combine both components. Furthermore, there are considerations on the meta-level, which, although subjective, strengthen the CaSc, e.g. considering a mouse model improper if it obviously demonstrates a full loss of function (knock-out) while the identified variant is a missense and highly suggestive for a gain of function.

Due to our awareness of the partial subjectivity of the CaSc, we aimed at proving its reproducibility. Evaluating the same variant by different scientists did not lead to deviating scoring, thus proving reproducibility of the CaSc. Certainly, it is important that people are well instructed and trained. It is also still not proven that the reproducibility holds up when the CaSC is used at different centers. We are aware that CaSc does not have a clear cut-off and that a high score is not a guarantee for a gene valid gene. We are also aware that CaSc is only a snapshot of supporting evidence at the time of analysis. However, our experience with published genes and ongoing studies as well as the reproducibility of CaSc show that it is an enormously useful tool in order to prioritize genes and to compare and communicate the relevance of candidate genes within the scientific community.

In opposite to the large studies, our approach does not allow making general statements on the genetics of NDD. However, our approach evaluates all cases, case by case, including the full spectrum of available clinical and genetic information. Case by case analysis seems at the first glance as a tedious work that cannot be done for a large number of affected individuals. However, after clinical examinations, often including imaging, followed by plenty of documentation tasks, wet lab sequencing, and bioinformatic preparation, the evaluation in a research setting of a trio exome and describing and documenting candidate genes represents only a small additional task but enriches the outcome enormously. Our experience shows that a trained scientist can easily handle several hundred cases per year, and that CaSc is easy to learn and can practically be implemented in daily clinical and research routine. Thus, we consider this scientific effort as feasible and necessary. Indeed, we consider it unjustifiable not to perform this last and rather small step after enormous efforts have been invested in each case.

As an outlook, large parts of CaSc can be automatized. However, at the time we see a big benefit in the manual evaluation of the literature. Future studies may compare the usability of the score in its current manual form and in an automatized form to see if there is a significant difference in performance. Also, such scoring systems can be expanded to other phenotypes, after proper modifications.

## Conclusions

In conclusion, our experience shows that case-specific evaluation of exome sequencing data followed by standardized scoring of candidate genes is a powerful first step to identify novel NDD genes. Probably tens of thousands of trio exome sequencing of NDD would benefit from such a tool since it would ease comparisons and communications, help scientists to make the most use of their results and speed deciphering of NDD.

## Supporting information

Supplementary Table 1 Info on variants

Supplementary Table 2 Info on patients

## Declarations

### Ethics approval and consent to participate

All analyses were performed in concordance to the provisions of the German Gene Diagnostic Act (*Gendiagnostikgesetz)* and the German Data Protection Act (*Bundesdatenschutzgesetz).* The project was approved by the ethic committee of the University of Leipzig, Germany (224/16-ek and 402/16-ek). Written informed consent of all examined individuals or their legal representatives was obtained prior to genetic testing and after advice and information about the risks and benefits of the study. This study does not involve animals.

### Consent for publication

Our manuscript does not contain any individual person’s data in any form (including any individual details, images or videos). As mentioned above, consent for publication has been obtained from every person, or in the case of children, their parent or legal guardian.

### Availability of data and material

The datasets generated and analyzed during the current study are available from the corresponding author Rami Abou Jamra on reasonable request. All results during this study are included in this published article and its supplementary information files.

### Competing interests

Ben Liesfeld and Roland Ewald work for Limbus and own shares. Saskia Biskup works for CeGat and owns shares. Peter Bauer works for Centogene. Dagmar Huhle works for Praxis für Humangenetik Leipzig and owns shares. All other authors declare that they have no competing interests.

### Funding

This study was not supported by funding.

### Authors’ contributions

BB, SM, AF, CBB, CH, JH, SuH, DLD, KP and RAJ have analyzed the data. BB and RAJ have written the manuscript. BB, MZ, JRL, DLD, KP and RAJ have contributed to concept and design. TB, AB, MKB, NDD, ME, CH, YH, SaH, FH, DH, SBK, WK, IK, AK, AM, DM, PM, RP, TP, IS, JUS, SS, DW, and MZ have examined and counseled patients. BL, RE, SB, PP and JH have generated data.

All authors have approved the submitted version. All authors have agreed both to be personally accountable for the author’s own contributions and to ensure that questions related to the accuracy or integrity of any part of the work, even ones in which the author was not personally involved, are appropriately investigated, resolved, and the resolution documented in the literature.

#### Acknowledgments

We thank all patients and their families who participated in this study.

### Web links and URLs

Genome Aggregation Database (GnomAD), https://gnomad.broadinstitute.org/

GeneMatcher, https://genematcher.org

Online Mendelian Inheritance in Man (OMIM), http://www.ncbi.nlm.nih.gov/Omim/

Sequence Ontology, http://www.sequenceontology.org/

VARVIS (Limbus, Rostock), https://www.limbus-medtec.com/

Genotype-Tissue Expression Project, https://gtexportal.org

## Supplemental Data

Supplemental Data includes four tables

**1. Supplemental Table 1 gives a good overview of the examined cases:** we <u>list all examined persons</u> sequentially from 1 to 198. We include general and clinical information on each individual, if the case is clinically solved (including gene), if we have identified a candidate gene or several candidate genes, or if the case remained fully negative. We contribute also information on the testing that has been performed (panel, exome, trio, single, etc.). We offer the table as pdf and as Excel files. The latter is easier to navigate in.

**2. Supplemental Table 2 gives a detailed description of identified genes and variants:** we <u>list all genes</u> in which we have identified variants. This includes candidate genes and also well-established NDD genes. We also list, to be complete, all fully negative cases. We add all available information on the clinical aspects, age, sex, family history, as well as all available information on the variant, the gene, the scoring and the rationale for our decision. We recommend using the Excel file in order to navigate in this table.

**3. Supplemntal Table 3 shows how different users score the same variant and gene:** If you are interested to know how people differ in scoring, check this table. Otherwise, it does not include much information.

**4. Supplemental Table** 4 **describes in details the scoring system:** If you want to implement scoring at your lab, you need this table in order to see our elaboration on how and when to score variants and genes.

