## Supplementary Table 3 for "Scientific evaluation of negative exome sequencing followed by systematic scoring of candidate genes to decipher the genetics of neurodevelopmental disorders"

| | CaSc total | | | | | | | | | | | | | Expression | | | | Hints to be involved in neuronal function | | | | Animal model with neuronal phenotype | | | | Hits | | | | related genes, interactions | | | |
|---|---|---|---|---|---|---|---|---|---|---|---|---|---|---|---|---|---|---|---|---|---|---|---|---|---|---|---|---|---|---|---|---|---|
| HGNC ID of candidate gene | BB | KP | AF | BB-KP | Difference BB-KP | BB-AF | Difference BB-AF | KP-AF | Difference KP AF | Average (BB,KP,AF) | standard deviation (BB,KP,AF) | BB-MW | Difference BB im Vgl MW | expression BB | expression KP | expression AF | standard deviation | hints neuronal function BB | hints neuronal function KP | hints neuronal function AF | standard deviation | animal models BB | animal models KP | animal models AF | standard deviation | hits BB | hits KP | hits AF | standard deviation | related genes, interactions BB | related genes, interactions KP | related genes, interactions AF | standard deviation |
| TTC28 | 7.7 | 7.0 | 6.7 | 0.7 | 0.7 | 1.0 | 1.0 | 0.3 | 0.3 | 7.1 | 0.4 | 0.6 | 0.6 | 0.4 | 0.4 | 0.4 | 0.00 | 0.5 | 0 | 0.5 | 0.24 | 0 | 0.5 | 0 | 0.24 | 1.32 | 0.66 | 0.33 | 0.41 | 0 | 0 | 0 | 0.00 |
| TEX13C | 4.0 | 4.0 | 3.5 | 0.0 | 0.0 | 0.5 | 0.5 | 0.5 | 0.5 | 3.8 | 0.2 | 0.2 | 0.2 | 0 | 0 | 0 | 0.00 | 0 | 0 | 0 | 0.00 | 0 | 0 | 0 | 0.00 | 0 | 0 | 0 | 0.00 | 0 | 0 | 0 | 0.00 |
| PLXNA3 | 5.8 | 6.5 | 6.8 | -0.7 | 0.7 | -1.0 | 1.0 | -0.3 | 0.3 | 6.4 | 0.4 | -0.5 | 0.5 | 0.7 | 0.7 | 0.7 | 0.00 | 1 | 1 | 1 | 0.00 | 0.5 | 0.5 | 0.5 | 0.00 | 0.33 | 1 | 1.32 | 0.41 | 1 | 1 | 1 | 0.00 |
| GRIN3B | 9.9 | 10.4 | 8.5 | -0.5 | 0.5 | 1.4 | 1.4 | 1.9 | 1.9 | 9.6 | 0.8 | 0.3 | 0.3 | 0.4 | 0.4 | 0.7 | 0.14 | 1 | 1 | 1 | 0.00 | 0.5 | 1 | 0.5 | 0.24 | 0 | 0 | 0.66 | 0.31 | 1 | 1 | 1 | 0.00 |
| NPTX1 | 9.1 | 8.1 | 8.6 | 1.0 | 1.0 | 0.5 | 0.5 | -0.5 | 0.5 | 8.6 | 0.4 | 0.5 | 0.5 | 1 | 1 | 1 | 0.00 | 1 | 0.5 | 1 | 0.24 | 1 | 0.5 | 0.5 | 0.24 | 0 | 0 | 0 | 0.00 | 1 | 1 | 1 | 0.00 |
| HIST1H2BC | 4.8 | 4.8 | 5.5 | 0.0 | 0.0 | -0.7 | 0.7 | -0.7 | 0.7 | 5.0 | 0.3 | -0.2 | 0.2 | 0 | 0 | 0.4 | 0.19 | 0 | 0 | 0 | 0.00 | 0 | 0 | 0 | 0.00 | 0 | 0 | 0 | 0.00 | 0 | 0 | 0 | 0.00 |
| GRIA4 | 9.3 | 9.0 | 9.3 | 0.3 | 0.3 | 0.0 | 0.0 | -0.3 | 0.3 | 9.2 | 0.2 | 0.1 | 0.1 | 1 | 1 | 1 | 0.00 | 1 | 1 | 1 | 0.00 | 1 | 1 | 1 | 0.00 | 0.33 | 0 | 0.33 | 0.16 | 1 | 1 | 1 | 0.00 |
| DUX4L4 | 7.0 | 7.0 | 6.5 | 0.0 | 0.0 | 0.5 | 0.5 | 0.5 | 0.5 | 6.8 | 0.2 | 0.2 | 0.2 | 0 | 0 | 0 | 0.00 | 0 | 0 | 0 | 0.00 | 0 | 0 | 0 | 0.00 | 0 | 0 | 0 | 0.00 | 0 | 0 | 0 | 0.00 |
| ZNF888 | 7.0 | 7.0 | 7.4 | 0.0 | 0.0 | -0.4 | 0.4 | -0.4 | 0.4 | 7.1 | 0.2 | -0.1 | 0.1 | 0 | 0 | 0.4 | 0.19 | 0 | 0 | 0 | 0.00 | 0 | 0 | 0 | 0.00 | 0 | 0 | 0 | 0.00 | 0 | 0 | 0 | 0.00 |
| FAM83G | 2.6 | 1.6 | 2.9 | 1.0 | 1.0 | -0.3 | 0.3 | -1.3 | 1.3 | 2.4 | 0.6 | 0.2 | 0.2 | 0 | 0 | 0 | 0.00 | 0 | 0 | 0 | 0.00 | 0 | 0 | 0 | 0.00 | 0 | 0 | 0.33 | 0.16 | 1 | 0 | 1 | 0.47 |
| NLRX1 | 4.4 | 5.3 | 4.4 | -0.9 | 0.9 | 0.0 | 0.0 | 0.9 | 0.9 | 4.7 | 0.4 | -0.3 | 0.3 | 0.4 | 0.4 | 0.4 | 0.00 | 0 | 0.5 | 0 | 0.24 | 0 | 0 | 0 | 0.00 | 0.33 | 0.66 | 0.33 | 0.16 | 0 | 0 | 0 | 0.00 |
| POLR1B | 5.6 | 5.9 | 5.6 | -0.3 | 0.3 | 0.0 | 0.0 | 0.3 | 0.3 | 5.7 | 0.1 | -0.1 | 0.1 | 0.4 | 0.7 | 0.4 | 0.14 | 0 | 0 | 0 | 0.00 | 0 | 0 | 0 | 0.00 | 0 | 0 | 0 | 0.00 | 0 | 0 | 0 | 0.00 |
| HIST1H4B | 4.2 | 4.9 | 4.2 | -0.7 | 0.7 | 0.0 | 0.0 | 0.7 | 0.7 | 4.4 | 0.3 | -0.2 | 0.2 | 0 | 0.7 | 0 | 0.33 | 0 | 0 | 0 | 0.00 | 0 | 0 | 0 | 0.00 | 0 | 0 | 0 | 0.00 | 0 | 0 | 0 | 0.00 |
| PPP1R37 | 7.2 | 5.9 | 6.5 | 1.3 | 1.3 | 0.7 | 0.7 | -0.6 | 0.6 | 6.5 | 0.5 | 0.7 | 0.7 | 0.7 | 0.7 | 0.7 | 0.00 | 0 | 0 | 0 | 0.00 | 0 | 0 | 0 | 0.00 | 0.66 | 0.33 | 0 | 0.27 | 1 | 0 | 1 | 0.47 |
| AQP6 | 5.4 | 4.9 | 5.8 | 0.5 | 0.5 | -0.4 | 0.4 | -0.9 | 0.9 | 5.4 | 0.4 | 0.0 | 0.0 | 0 | 0 | 0.4 | 0.19 | 0.5 | 0 | 0.5 | 0.24 | 0 | 0 | 0 | 0.00 | 0 | 0 | 0 | 0.00 | 0 | 0 | 0 | 0.00 |
| PSMB3 | 5.4 | 5.1 | 4.7 | 0.3 | 0.3 | 0.7 | 0.7 | 0.4 | 0.4 | 5.1 | 0.3 | 0.4 | 0.4 | 0.7 | 0.4 | 0 | 0.29 | 0 | 0 | 0 | 0.00 | 0 | 0 | 0 | 0.00 | 0 | 0 | 0 | 0.00 | 0 | 0 | 0 | 0.00 |
| KANK4 | 6.9 | 6.9 | 6.8 | 0.0 | 0.0 | 0.1 | 0.1 | 0.1 | 0.1 | 6.9 | 0.0 | 0.0 | 0.0 | 0.4 | 0.4 | 0.4 | 0.00 | 0 | 0 | 0 | 0.00 | 0 | 0 | 0 | 0.00 | 0 | 0 | 0 | 0.00 | 0 | 0 | 0 | 0.00 |
| LCTL | 6.9 | 7.2 | 7.7 | -0.3 | 0.3 | -0.8 | 0.8 | -0.5 | 0.5 | 7.3 | 0.3 | -0.4 | 0.4 | 0.4 | 0.4 | 0.7 | 0.14 | 0 | 0 | 0 | 0.00 | 0 | 0 | 0 | 0.00 | 0 | 0.33 | 0 | 0.16 | 0 | 0 | 0 | 0.00 |
| TOB1 | 9.2 | 10.2 | 9.2 | -1.0 | 1.0 | 0.0 | 0.0 | 1.0 | 1.0 | 9.5 | 0.5 | -0.3 | 0.3 | 0.7 | 0.4 | 0.7 | 0.14 | 0.5 | 0.5 | 0.5 | 0.00 | 0 | 0 | 0 | 0.00 | 0 | 0.33 | 0 | 0.16 | 0 | 1 | 0 | 0.47 |
| ACTL6B | 10.0 | 10.8 | 10.5 | -0.8 | 0.8 | -0.5 | 0.5 | 0.3 | 0.3 | 10.4 | 0.3 | -0.4 | 0.4 | 1 | 1 | 1 | 0.00 | 1 | 1 | 1 | 0.00 | 0.5 | 1 | 1 | 0.24 | 1.32 | 1.66 | 1.32 | 0.16 | 1 | 1 | 1 | 0.00 |
| CELSR3 | 6.6 | 6.8 | 6.4 | -0.2 | 0.2 | 0.2 | 0.2 | 0.4 | 0.4 | 6.6 | 0.2 | 0.0 | 0.0 | 1 | 1 | 1 | 0.00 | 1 | 1 | 0.5 | 0.24 | 0.5 | 1 | 0.5 | 0.24 | 0.66 | 0.33 | 0.66 | 0.16 | 1 | 1 | 1 | 0.00 |
| GPKOW | 2.5 | 3.2 | 3.7 | -0.7 | 0.7 | -1.2 | 1.2 | -0.5 | 0.5 | 3.1 | 0.5 | -0.6 | 0.6 | 0.7 | 0.7 | 0.7 | 0.00 | 0 | 0 | 0.5 | 0.24 | 0 | 0 | 0 | 0.00 | 0 | 0.66 | 0.66 | 0.31 | 0 | 0 | 0 | 0.00 |
| PABPC1 | 6.4 | 7.1 | 6.4 | -0.7 | 0.7 | 0.0 | 0.0 | 0.7 | 0.7 | 6.6 | 0.3 | -0.2 | 0.2 | 0.4 | 0.4 | 0.4 | 0.00 | 0 | 0 | 0 | 0.00 | 0 | 0 | 0 | 0.00 | 0.33 | 0 | 0.33 | 0.16 | 0 | 1 | 0 | 0.47 |
| CASKIN1 | 8.2 | 8.2 | 7.2 | 0.0 | 0.0 | 1.0 | 1.0 | 1.0 | 1.0 | 7.9 | 0.5 | 0.4 | 0.4 | 1 | 1 | 1 | 0.00 | 1 | 1 | 1 | 0.00 | 0.5 | 0.5 | 0.5 | 0.00 | 0 | 0 | 0 | 0.00 | 1 | 1 | 0 | 0.47 |
| FAM181B | 5.0 | 5.0 | 6.5 | 0.0 | 0.0 | -1.5 | 1.5 | -1.5 | 1.5 | 5.5 | 0.7 | -0.5 | 0.5 | 1 | 1 | 1 | 0.00 | 1 | 1 | 1 | 0.00 | 0 | 0 | 1 | 0.47 | 0 | 0 | 0 | 0.00 | 0 | 0 | 0 | 0.00 |
| DNAH17 | 2.3 | 2.6 | 4.3 | -0.3 | 0.3 | -2.0 | 2.0 | -1.7 | 1.7 | 3.1 | 0.9 | -0.7 | 0.7 | 0.4 | 0.7 | 0.7 | 0.14 | 0 | 0 | 0 | 0.00 | 0 | 0 | 0 | 0.00 | 0 | 0 | 0.66 | 0.31 | 0 | 0 | 1 | 0.47 |
| HOOK2 | 4.4 | 5.4 | 4.4 | -1.0 | 1.0 | 0.0 | 0.0 | 1.0 | 1.0 | 4.7 | 0.5 | -0.3 | 0.3 | 0.4 | 0.4 | 0.4 | 0.00 | 0 | 0.5 | 0 | 0.24 | 0 | 0 | 0 | 0.00 | 0 | 0 | 0 | 0.00 | 0 | 0.5 | 0 | 0.24 |
| KMT2E | 10.7 | 11.2 | 9.8 | -0.5 | 0.5 | 0.9 | 0.9 | 1.4 | 1.4 | 10.6 | 0.6 | 0.1 | 0.1 | 0.7 | 0.7 | 0.7 | 0.00 | 0 | 0.5 | 0 | 0.24 | 0 | 0 | 0 | 0.00 | 0.99 | 0.99 | 0.66 | 0.16 | 1 | 1 | 1 | 0.00 |
| SPEN | 9.4 | 10.3 | 8.9 | -0.9 | 0.9 | 0.5 | 0.5 | 1.4 | 1.4 | 9.5 | 0.6 | -0.1 | 0.1 | 0.7 | 0.4 | 0.7 | 0.14 | 0.5 | 1 | 0.5 | 0.24 | 1 | 1 | 0.5 | 0.24 | 1.32 | 2 | 1.32 | 0.32 | 1 | 1 | 1 | 0.00 |
| average | | | | | 0.50 | | 0.58 | | 0.76 | | 0.40 | | 0.30 | | | | 0.07 | | | | 0.07 | | | | 0.07 | | | | 0.13 | | | | 0.11 |
| standard deviation | | | | | 0.38 | | 0.51 | | 0.47 | | 0.19 | | 0.20 | | | | 0.10 | | | | 0.11 | | | | 0.12 | | | | 0.14 | | | | 0.19 |
