## Supplementary Table 4 for "Scientific evaluation of negative exome sequencing followed by systematic scoring of candidate genes to decipher the genetics of neurodevelopmental disorders"

### Table S4; Detailed description of the Candidate Score (CaSc)

**Inheritance**

| | | | |
|---|---|---|---|
| 1 | **zygosity/family history/segregation** | *de novo* | 2 |
| | There is between 1 and 2 *de novo* variants per exome. In an outbred family, *de novo* is the most common cause of NDD. Thus, if there is a *de novo* variant, it automatically receives a high scoring of 2. Also of relevance are homozygous variants since these are rare in outbred families and we thus give 2 points. Also we boost scoring for cases with an affected male and a second, maternal, affected male if the variant is on the chromosome X. Less impressive are compound heterozygous variants since our experience shows that practically everybody has 1-5 rare, compound heterozygous variants. The same accounts for hemizygous X chromosomal variants. Most convincing is a bi-allelic variant (homo or compound heterozygous) in a family that most probably has an autosomal recessive phenotype (more than one affected child) and thus gets a score of 3. | homo or comphet with ≥ 2 affected children | 3 |
| | | homo | 2 |
| | | comphet | 1 |
| | | X linked and a boy | 1 |
| | | X linked and at least a second affected maternal male relative | 2 |
| | | other | 0 |

**Gene attributes**

| | | | |
|---|---|---|---|
| 2 | **pLI-score** | <0.9 | 0 |
| | This parameter describes if a gene is possibly intolerant for heterozygous loss of function (LoF). We use the pLI score of GnomAD (but using other scores such as RVIS may be as good) to reflect intolerance of LoF. We decided to give 1 point to genes with pLI-score ≥0.9 since these are probably loss-of-function intolerant as suggested by the authors of pLI. The rest get no points. | ≥0.9 | 1 |
| 3 | **missense z-score** | < 0 | 0 |
| | For missense variants we aimed at a similar approach compared to the pLI-score. z score of ~3.09 is equivalent to a p-value of $10^{-3}$ and is considered the significance threshold when splitting transcripts into constrained and unconstrained classes Thus, we decided to give one point to a z-score of ≥ 3.09, no points for negative z-scores and the rest get 0.5 points. | 0-3.08 | 0.5 |
| | | ≥3.09 | 1 |
| 4 | **expression** | not in CNS | 0 |
| | We have used for the analyses described in this manuscript the Genotype-Tissue Expression (GTEx) Project. GTEx Portal: https://gtexportal.org. Here we want to reflect the expression of the gene in the central nervous system (CNS) comparable to other tissues. It would have been theoretically possible to make this point fully objective by downloading expression data and running comparisons between brain and other tissues expression. However, we realized that this would lead to complex scoring with a risk of less compliance. So we decided to leave this parameter partially subjective and we did not make clear cuts of expression. One point if a gene is exclusively expressed in brain, 0 if it is overall not expressed in brain. For genes in between these extremes, we went for 0.4 and 0.7 as described to the right. | low in CNS, and more in (some) other tissues | 0.4 |
| | | expressed in CNS and is comparable to (some) other tissues | 0.7 |
| | | most or exclusively in CNS | 1 |

| | | | |
|---|---|---|---|
| **Variant attributes** | | | |
| 5 | **assumed impact on protein**<br><br>To estimate the impact on protein, we used the categories High, Moderate, and Low as described by the Sequence Ontology (SO) terms (http://www.sequenceontology.org/). Basically, however, truncating variants would be high while missense variants as well as splice variants that are not at the consensus splice site are moderate. | moderate | 0 |
| | | high and heterozygous | 2 |
| | | high and bi-allelic | 3 |
| | | comphet = one moderate and one high | 1 |
| | | other | 0 |
| 6 | ***in silico* parameters**<br><br>Practically all variants that are "high" based on SO gets 1 point. In order to evaluate missense variants adequately, we implemented the *in silico* evaluation of the variants. For several *in silico* prediction programs we spotted the value on a percentile scale from 0 to 1 (so called "ranking" whereas 0 is benign and 1 is pathogenic) and added the average of available *in silico* programs (preferably Sift, MutationTaster, and Mutation Assessor) to the score. We do not have the percentile parameters for splicing variants, thus, if two or more splicing prediction programs (we used SpliceSiteFinder-like, MaxEntScan, NNSPLICE, GeneSplicer, and Human Splicing Finder) were positive, we consider the variant "high". If there are no available *in silico* parameters, we allocate the median value of these parameters, i.e. 0.5 points, the same if only one splicing program reveal a splicing effect. | missense[e] | average |
| | | LoF | 1 |
| | | splicing affected in one program[f] | 0.5 |
| | | splicing affected in two or more programs[f] | 1 |
| | | no available in silico values | 0.5 |
| 7 | **conservation**<br><br>We used GERP++RS for the estimation of conservation and estimated this based on other parameters if this value was not available. For truncating variants the conservation is irrelevant, thus it gets 1 point. | LoF | 1 |
| | | percentile/ranking of the values of all possible variants[g] | 0-1 |
| 8 | **frequency**<br><br>We have used the GnomAD, 1000 genome project as we all our internal database of about 3000 probes. For autosomal dominant NDD phenotypes, we expect a minor allele frequency (MAF) of 0. If this is achieved, the variant gets 1 point. However, we have left an error range (due to artefacts or exceptions) of one allele to be identified in all above mentioned databases together. If the phenotype of the patient is rather mild and we think that reproduction is possible, we allow a higher MAF of up to 5 alleles. For the autosomal recessive genes, we assumed that ultra-rare variants (MAF < 0.00005 in GnomAD) would get a higher scoring than more common variants (MAF < 0.0005) as described to the right. For variants on chromosome X we assumed for severe phenotypes that a discrepancy between females and males (i.e. there are some alleles in females and none or only one (as an exception due to an artefact) is striking and gets 2 points. | *de novo* or inherited in an AD pattern MAF of 0 or a maximum of one allele in all available databases in a disorder with no or extremely low reproduction chance | 1 |
| | | *de novo* or inherited in an AD pattern in a disorder with a chance for reproduction;<br><br>max. allele number of 5 (MAF ≈ 0.00002) in GnomAD | 0.5 |
| | | autosomal recessive inheritance; max. MAF of 0.0005 in GnomAD[i] | 0.5 |
| | | autosomal recessive inheritance; MAF of max 0.00005[i] | 1 |

| | | | |
|---|---|---|---|
| | | X linked and discrepancy of MAF in GnomAD between males and females in a disorder with no chance for reproduction[j] | 2 |
| | | other | 0 |

**Literature research**

| | | | |
|---|---|---|---|
| 9 | **neuronal function**<br><br>If there is no evidence in the scientific literature for a neuronal function of the gene or gene product we allocate 0 points. If there is evidence in the scientific literature for unspecific neuronal functions (regarding the evaluated phenotype of NDD) of the gene or gene product we allocate 0.5 points. If there is evidence in scientific literature for specific neuronal functions of the gene or gene product similar to the evaluated phenotype we allocate 1 point. E. g. for genes with 1 point are these with functions in signal transduction or development of the CNS. The information on the functions are obtained from PubMed and is prone for subjective evaluation in dependence of the scientist who perform the scoring. Making this parameter objective is difficult and would make scoring much more complex. We thus decided to take the risk of having some subjectivity. | no hints to be involved in neuronal function | 0 |
| | | hints to be involved in neuronal function | 0.5 |
| | | hints to be involved in neuronal function regarding signaling/development | 1 |
| 10 | **gene family/neurological interactions**<br><br>This parameter is very much similar to the parameter above and also has very much overlapping with the parameter below. We decided to divide the "Literature Research" to these three parameters to reduce the effect of a failure. All three parameters are subjective to some extent and can actually be bundled in one large parameter of up to 3 or even 4 points. But then, a mistake in scoring would have large influence. While dividing the parameters "literature" in three smaller ones leads to less serious consequences of mistakes. | yes | 1 |
| | | no | 0 |
| 11 | **animal model**<br><br>If there is evidence in the scientific literature for animal models of the candidate gene with neurological or behavioral phenotypes we allocate 0.5 points. If the phenotype of the model overlaps with aspects of NDD, we allocate 1 point. Otherwise, no points. | none | 0 |
| | | no neurological or behavioral phenotype | 0 |
| | | neurological or behavioral phenotype | 0.5 |
| | | comparable neurological phenotypes | 1 |
| 12 | **reported somewhere else as candidate for NDD?**<br><br>There are many lists and tables with reported candidate genes/variants in the literature. Also, we have internal lists of candidate genes/variants that we received from cooperation partners. If the candidate gene was reported in other studies with neurological or comparable phenotype, we allocate 0.33 points per hit. To restrain the power of this category, we limit the max. score to 2 points (i.e. 6 hits). When performing this scoring, we consider the quality of the reference and use this category restrictively. Also here, there is a possibility to reduce subjectivity by pre-defining the sources to use. However, in our opinion the loss of information would be an even greater disadvantage compared to using all | per hit, *max. score: 2* | 0.33 |

individual available sources of information. We usually use the sources HGMD, ClinVar, PubMed search, and occasionally GeneMatcher (GeneMatcher is rather used if a gene already achieve a good scoring). It is clear that this aspect cannot be exhaustive.
